# Antimicrobial activity of a repurposed harmine-derived compound on extensively drug-resistant *Acinetobacter baumannii* clinical isolates

**DOI:** 10.1101/2021.06.30.450460

**Authors:** Anke Breine, Mégane Van Gysel, Mathias Elsocht, Clémence Whiteway, Chantal Philippe, Théo Quinet, Adam Valcek, Johan Wouters, Steven Ballet, Charles Van der Henst

**Affiliations:** Microbial Resistance and Drug Discovery, VIB-VUB Center for Structural Biology, VIB, Flanders Institute for Biotechnology, Brussels, Belgium; Structural Biology Brussels, Vrije Universiteit Brussel (VUB), Brussels, Belgium; Namur Medicine & Drug Innovation Center (NAMEDIC), University of Namur (UNamur), Namur, Belgium; Research Group of Organic Chemistry, Departments of Chemistry and Bioengineering Sciences, Vrije Universiteit Brussel, Pleinlaan 2, B-1050 Brussels, Belgium; Research Unit in the Biology of Microorganisms (URBM), NARILIS, University of Namur (UNamur), Belgium; Molecular Biology and Evolution, C.P. 160/16 Université Libre de Bruxelles (ULB), Avenue F.D. Roosevelt 50, B-1050 Brussels, Belgium

## Abstract

**Objectives:** The spread of antibiotic resistant bacteria is an important threat for human healthcare. *Acinetobacter baumannii* bacteria impose one of the major issues, as multidrug- to pandrug-resistant strains have been found, rendering some infections untreatable. In addition, *A. baumannii* is a champion in surviving in harsh environments, being capable of resisting to disinfectants and to persist prolonged periods of desiccation. Due to the high degree of variability found in *A. baumannii* isolates, the search for new antibacterials is challenging. Here, we screened a compound library to identify compounds active against recent isolates of *A. baumannii* bacteria.

**Methods:** A repurposing drug screen was undertaken to identify *A. baumannii* growth inhibitors. One hit was further characterized by determining its IC_50_ and testing its activity on 43 recent clinical *A. baumannii* isolates, amongst which 40 are extensively drug- and carbapenem-resistant strains.

**Results:** The repurposing screen led to the identification of a harmine-derived compound, called HDC1, which proved to have bactericidal activity on the multidrug-resistant AB5075-VUB reference strain with an IC_50_ of 48.23 µM. In addition, HDC1 impairs growth of all 43 recent clinical *A. baumannii* isolates.

**Conclusions:** We identified a compound with inhibitory activity on all tested, extensively drug-resistant clinical *A. baumannii* isolates.

## Introduction

The rise of antibiotic resistant bacteria is a global threat for healthcare, making it possible to succumb to diseases that were previously treatable. ^1^ This has been acknowledged by both the World Health Organization (WHO) and the Centers for Disease Control and Prevention (CDC), which generated a list of drug-resistant pathogens for which new antibiotics are urgently needed. ^2,3^ The top priorities of these lists are antibiotic-resistant *Acinetobacter baumannii* bacteria.

*A. baumannii* is a Gram-negative, opportunistic bacterium, belonging to the ESKAPE group (*Enterococcus faecium, Staphylococcus aureus, Klebsiella pneumoniae, Acinetobacter baumannii, Pseudomonas aeruginosa* and *Enterobacter* spp.) of nosocomial pathogens. ^4,5^ While the pathogen is ubiquist (*i*.*e*. it can be found in soil, on human skin and in water sources), its presence especially imposes a threat in clinical settings. ^6,7^ This is due to a remarkable combination of resistance capabilities of *A. baumannii*, which is able to persist prolonged periods of desiccation, to resist to disinfectants and to acquire drug resistance at a high rate. ^8,9^ Infections caused by *A. baumannii* commonly occur in immunocompromised patients and manifest as ventilator-assisted pneumonia, bacteremia and to a lesser extent skin or urinary tract infections. ^10^ Treatment of these infections becomes increasingly difficult, as multidrug-resistant, to extensively drug-resistant or even pandrug-resistant strains have been reported, with the latter being resistant to all available antibiotics, including carbapenems. ^11,12^ An important hurdle in the development of new antimicrobials against *A. baumannii* is the high diversity found between isolates, leading to a still open pan-genome. ^13^

In this paper, we aimed at the discovery of a compound active against most clinical isolates. We performed a repurposing screen on a compound library, which led to the identification of a <u>h</u>armine-<u>d</u>erived <u>c</u>ompound, called HDC1, with inhibitory activity on the growth of all the tested recent clinical isolates, amongst which 40 are extensively drug- and carbapenem-resistant.

## Material and methods

### Compound library and synthesis of HDC1

A compound library of the Namur Medicine & Drug Innovation Center (NAMEDIC) was provided for a growth inhibition screen against *A. baumannii*. All compounds were dissolved in 100% DMSO and used for the initial screen at 100 µM. The active compound HDC1 (1-methyl-2-benzyl-7-benzyloxy-9-benzyl-β-carbolin-2-ium bromide) was synthesized as previously described ^14^ with the following optimizations: 1-methyl-7-hydroxy-β-carboline was synthesized by adding 1-methyl-7-methoxy-β-carboline (0.600 g, 2.83 mmol, 1 equiv.), hydrobromic acid (12 ml, 48% in H_2_O) and acetic acid (12 ml) into a round-bottom flask, equipped with reflux condenser. The mixture was refluxed overnight under argon atmosphere and subsequently added to distilled water (100 ml). The precipitate was isolated via filtration, washed with cold water and dried under vacuum yielding 1-methyl-7-hydroxy-β-carboline with 81% (0.454 g) yield. ^1^ H-NMR (500 MHz, DMSO-d6) δ (ppm): 12.59 (s, 1H), 10.62 (s, 1H), 8.39-8.29 (m, 2H), 8.21 (d, J = 8.7 Hz, 1H), 7.03 (d, J = 1.7 Hz, 1H), 6.89 (dd, J = 8.7, 1.7 Hz, 1H), 2.94 (s, 3H). Next, 1-methyl-2-benzyl-7-benzyloxy-9-benzyl-β-carbolin-2-ium bromide was synthesized. First 1-methyl-7-hydroxy-β-carboline (0.500 g, 2.52 mmol, 1 equiv.) was dissolved in anhydrous N,N,-dimethylformamide (20 ml) into a flame-dried microwave vial under argon atmosphere. Then KOtBu (0.849 g, 7.57 mmol, 3 equiv.) was added and the mixture was stirred for 30 minutes at room temperature. Subsequently benzyl bromide (3.00 ml, 25.2 mmol, 10 equiv.) was added and the mixture was heated overnight at 75°C. Afterwards the crude mixture was filtered, and the precipitate was washed with CH_2_Cl_2_. The volatiles in the filtrate were removed under reduced pressure and the crude product was subjected to column chromatography (cyclohexane/ethyl acetate) yielding the desired product with 46% (0.638 g) yield. ^1^ H-NMR (500 MHz, DMSO-d6) δ (ppm): 8.86 (d, J = 6.6 Hz, 1H), 8.72 (d, J = 6.6 Hz, 1H), 8.49 (d, J = 8.8 Hz, 1H), 7.57 (d, J = 2.1 Hz, 1H), 7.50-7.46 (m, 2H), 7.40-7.25 (m, 9H), 7.23 (dd, J = 8.8, 2.1 Hz, 1H), 7.12 (d, J = 7.2 Hz, 2H), 6.99 (d, J = 7.2 Hz, 2H), 6.02 (s, 2H), 5.99 (s, 2H), 5.27 (s, 2H), 2.85 (s, 3H). ^13^ C-NMR (126 MHz, DMSO-d6) δ (ppm): 162.8 (Cq), 148.0 (Cq), 139.6 (Cq), 137.5 (Cq), 136.2 (Cq), 135.4 (Cq), 134.7 (Cq), 133.5 (Cq), 129.1 (CH), 129.0 (CH), 128.5 (CH), 128.3 (CH), 127.5 (CH), 126.6 (CH), 125.4 (CH), 124.9 (CH), 114.7 (CH), 113.7 (CH), 112.8 (Cq), 95.0 (CH), 70.1 (CH2), 59.8 (CH2), 48.3 (CH2), 16.0 (CH3). The spectroscopic data were in accordance with those reported by Frédérick *et al*. (2012). ^14^

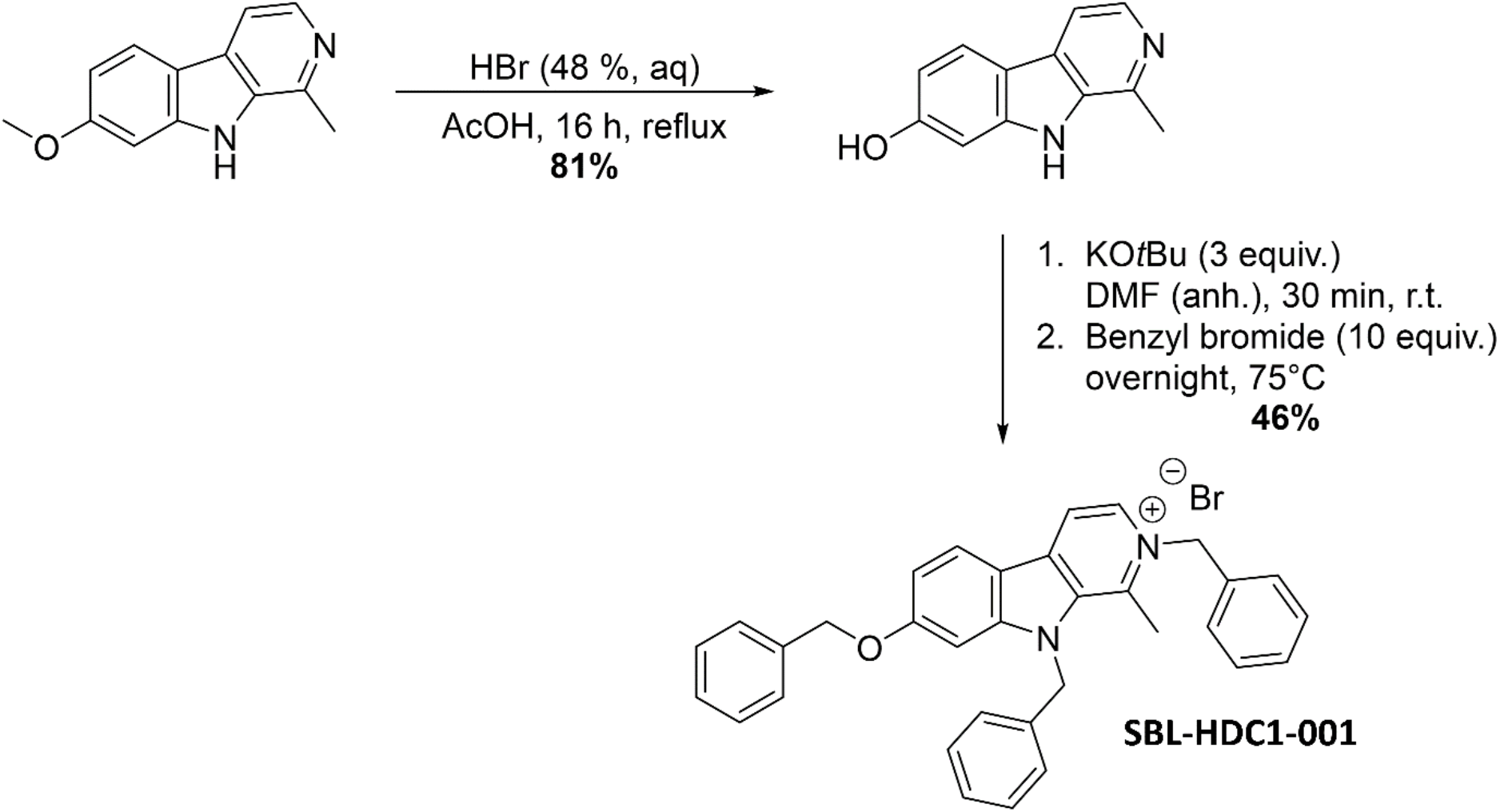

### OD measurements

All *A. baumannii* reference strains and clinical isolates were cultured at 37°C in LB broth media for 16h before subsequent experiments. For IC_50_ determination, bacterial cells corresponding to OD_600nm_=0.1 were transferred to a 96 well flat bottom plate (Greiner, Austria) containing varying concentrations of HDC1 (0.1; 1; 10; 25; 50; 75; 100 and 1000 µM) in LB broth media with 1% DMSO. For determination of the activity of HDC1 on the 43 clinical *A. baumannii* isolates, bacterial cells corresponding to OD_600nm_=0.1 were transferred to a 96 well flat bottom plate (Greiner, Austria) containing 100 µM of HDC1 in LB broth media with 1% DMSO. For both experiments, the positive control included bacterial cells corresponding to OD_600nm_=0.1 of the used strains in LB broth media with 1% DMSO. The negative control included both LB broth media and LB broth media supplemented with 1% DMSO. Collection of data was done using the Cytation 1 (BioTek, United States). The OD_600_ absorbance of all bacterial cultures was measured every 30 min for 24h at a temperature of 37°C and agitation of 355 cpm (cycles per minute). In all analysis, growth inhibition is defined as decreasing absorbance values over time compared to the positive control. The data were measured in biological triplicate.

### CFU determination

After absorbance measurements for IC_50_ determination, the positive control (AB5075-VUB grown in LB with 1% DMSO) and AB5075-VUB grown in presence of 100 µM HDC1, were resuspended in PBS, brought to the same OD_600nm_ and plated on LB agar plates in appropriate dilutions. After 16 h incubation of the LB agar plates at 37°C, CFUs were counted to assess any bacteriostatic or bactericidal effect. All data was measured in biological triplicate. To estimate the initial bacterial load, the relationship between the absorbance at OD_600nm_ and CFUs was determined for the strain AB5075-VUB. This was done by plating serial dilutions of bacterial cultures with a known OD_600nm_ on LB agar and counting the corresponding CFUs (see Table S1).

### IC_50_ calculation

For the determination of the IC_50_, GraphPad Prism 9 (GraphPad Software, LLC) was used. After 20 hours, the OD_600nm_ absorbance kinetic data were obtained and normalized using the positive control absorbance value as 100% viability and the absorbance value of 1 mM HDC1 as 0% viability. The analysis was then done on the normalized data by nonlinear regression curve fitting. The IC_50_ value is shown with a 95% confidence interval (CI).

### Statistical analysis

All data shown are represented as mean ± standard deviation of three biological replicates, except otherwise stated. CFU were statistically analyzed by an unpaired t-test. All growth curves were statistically analyzed by a Mann Whitney test. The p values < 0.05 were considered significant.

## Results

### Repurposing screen reveals compound with inhibitory activity on AB5075-VUB

The initial screen from a chemical library of the Namur Medicine & Drug Innovation Center from the University of Namur (UNamur) aimed at the identification of growth inhibitors for problematic multidrug-resistant *A. baumannii* strains. The strain initially used for this screen is a multidrug-resistant *A. baumannii* reference strain, AB5075-VUB. This strain is a derivative from the parental strain AB5075 that was clonally isolated in our laboratory at the VUB (Vrije Universiteit Brussel).

The screen showed complete growth inhibition by one compound called HDC1, for it is a <u>h</u>armine-<u>d</u>erived <u>c</u>ompound (**Figure 1**). Harmine is a natural -carboline known to have many pharmacological activities such as antimicrobial, antifungal and antitumor properties. ^14,15^ HDC1 was originally synthesized for an anticancer drug screen. Interestingly, compounds with anticancer activity have lately been explored as potential antimicrobials. ^16^ In line with this tendency, HDC1 was further characterized for other activities.

**Figure 1.**
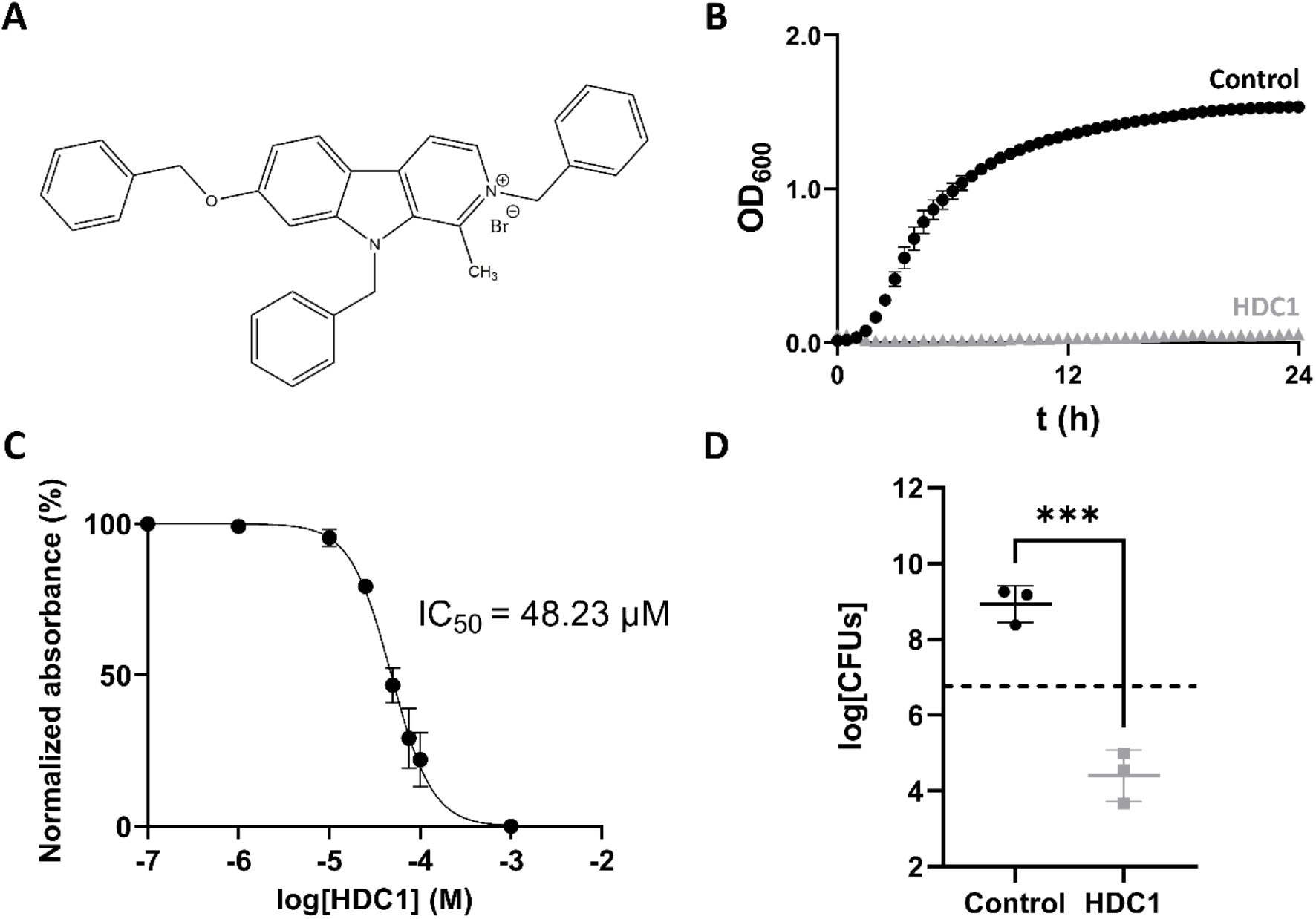
**A** Structure of HDC1. **B** Growth curve of AB5075-VUB in presence (grey triangles) and absence (black spheres) of HDC1. **C**. Non-linear regression curve of normalized absorbance reads in function of compound concentration. **D**. Number of viable bacteria after 24 h incubation without and with HDC1. The initial bacterial load is represented by the dashed line on the plot. All data points in this figure are shown as mean±standard deviation of three independent biological replicates.

### HDC1 has bactericidal activity on AB5075-VUB

To determine the potency of the compound, the minimum concentration required for 50% growth inhibition, IC_50_, was determined for the AB5075-VUB strain. The analysis showed that HDC1 has an IC_50_ of 48.23 µM (95% CI 44.76-51.83) (**Figure 1C**). To determine the potential antimicrobial effect of HDC1 on AB5075-VUB viability, the strain was grown in presence of 100 µM of HDC1. After 24 h incubation, the bacteria were resuspended in fresh media without the compound and plated on LB agar plates for CFUs enumeration. After 24h incubation, the number of recovered bacteria is significantly different when bacteria are incubated with the compound compared to the control group (**Figure 1D**). While an increase in CFUs is observed for the control group, a decrease of CFUs is observed in the presence of HDC1, compared to the initial bacterial load. This shows a significant bactericidal activity of HDC1 on the multidrug-resistant AB5075-VUB reference strain.

### HDC1 has broad inhibitory activity on all the tested clinical isolates

*A. baumannii* has a highly dynamic genome. ^17^ The presence of mobile genetic elements and the efficient acquisition of genes through horizontal gene transfer are not only responsible for the pathogen’s success in obtaining drug resistance and environmental persistence, but they are also the reason isolates have become more and more diverse. ^13,17,18^ The core genome of the pathogen’s strains is reported to be relatively small and the accessory genome of strains can be up to 25-46% unique. ^13,18^ This heterogeneity found in isolates renders the search for antimicrobial compounds increasingly difficult. It is therefore important for a new potential antimicrobial to exert its activity not only on a few *A. baumannii* strains, but on a multitude of diverse and clinically relevant isolates.

To determine the activity of HDC1 on recent isolates, we used 43 multidrug-resistant *A. baumannii* strains. These strains were isolated between 2014 to 2017, in different Belgian hospitals, of varying patient infections sites and have varying antibiograms. ^19^ The resistance profiles of 40 of these isolates show that they can even be classified as extensively drug-resistant, as previously defined for *Acinetobacter* spp. ^12^ In addition to these recent clinical isolates, three frequently used reference strains were also included. Two of these strains, ATCC19606 and ATCC17978, are older type strains, compared to the more recent and multidrug-resistant AB5075 reference strain. ^11,20^ The third reference strain, DSM30011, is an environmental isolate. ^21^

The growth of all tested strains was impaired by the presence of HDC1 (**Figure 2**), with low, intermediate, or complete inhibition levels. An overview of the different resistance profiles against HDC1 can be found in **Table 1**. Complete inhibition of growth is observed for all three reference strains and 17 recent isolates, while most of the clinical isolates show an intermediate inhibition profile. The AB193-VUB isolate showed the least sensitivity to HDC1. No correlation could be found between the sensitivity of the tested strains to HDC1 and the antibiogram profiles of the strains. ^19^

**Table 1.** Growth inhibition levels on three reference strains and 43 recent clinical isolates of *A. baumannii*

| Level of growth inhibition |  |  |
| --- | --- | --- |
| Low | Intermediate | Complete |
| AB193-VUB | AB3-VUB | ATCC19606 |
|  | AB9-VUB | ATCC17978 |
|  | AB14-VUB | DSM30011 |
|  | AB21-VUB | AB16-VUB |
|  | AB36-VUB | AB20-VUB |
|  | AB40-VUB | AB32-VUB |
|  | AB167-VUB | AB39-VUB |
|  | AB169-VUB | AB176-VUB |
|  | AB171-VUB | AB177-VUB |
|  | AB172-VUB | AB186-VUB |
|  | AB173-VUB | AB187-VUB |
|  | AB175-VUB | AB189-VUB |
|  | AB179-VUB | AB194-VUB |
|  | AB180-VUB | AB212-VUB |
|  | AB181-VUB | AB214-VUB |
|  | AB183-VUB | AB220-VUB |
|  | AB188-VUB | AB227-VUB |
|  | AB193-VUB | AB229-VUB |
|  | AB213-VUB | AB231-VUB |
|  | AB216-VUB | AB232-VUB |
|  | AB217-VUB |  |
|  | AB219-VUB |  |
|  | AB222-VUB |  |
|  | AB224-VUB |  |
|  | AB226-VUB |  |
|  | AB233-VUB |  |

**Figure 2.**
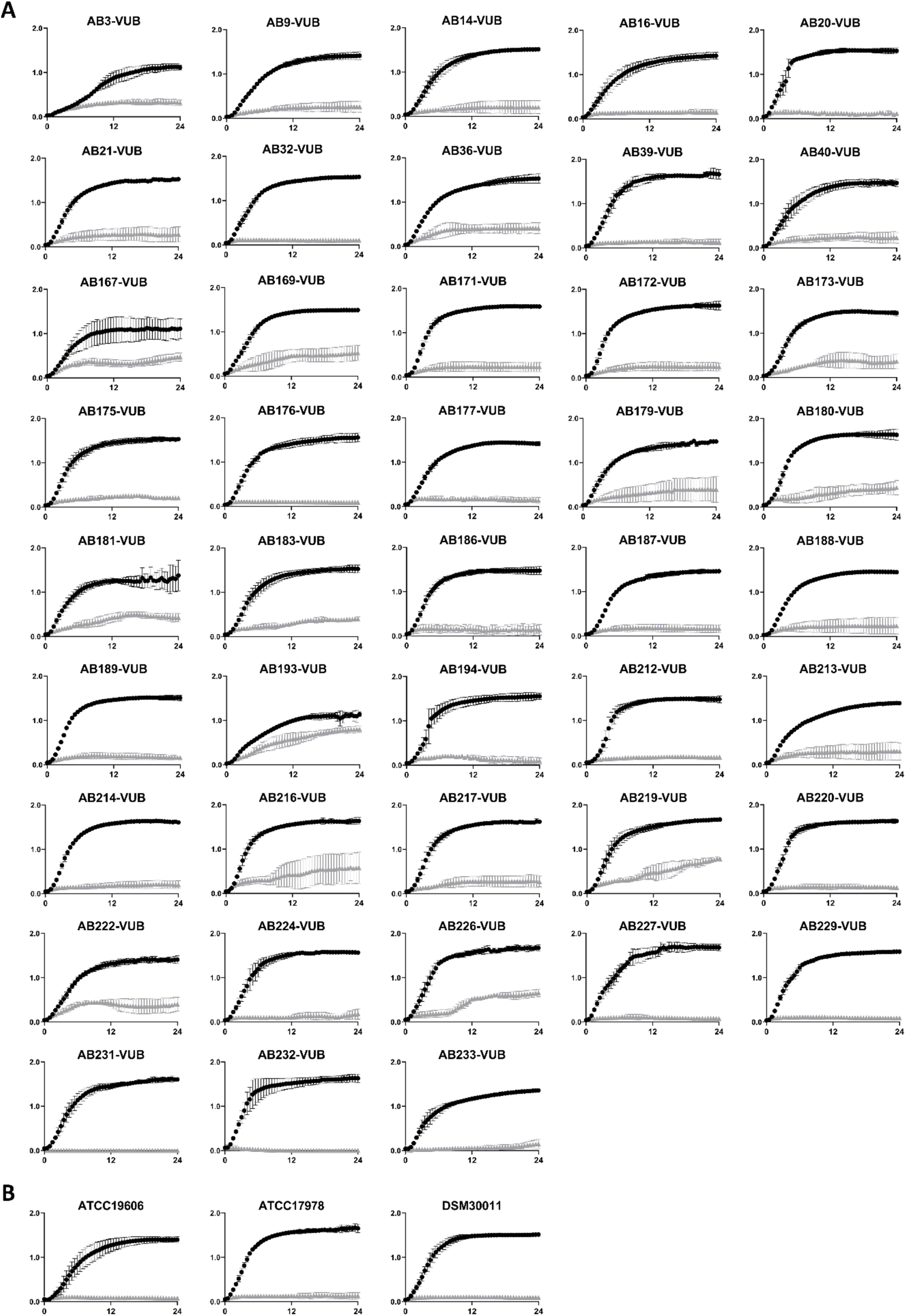
**A**. Growth curves of 43 recent clinical *A. baumannii* isolates. **B**. Growth curves of 3 *A. baumannii* reference strains (ATCC19606, ATCC17978 and DSM30011). All strains were incubated with 100 µM of HDC1 (grey triangles) or without HDC1 (black spheres) for 24 h. All data points are shown as mean standard deviation of three independent biological replicates. X axis and y axis correspond to respectively time (h) and absorbance values measured at 600 nm.

## Discussion

In this study, a repurposing drug screen led to the discovery of a compound with inhibitory activity on the growth of a multidrug-resistant *Acinetobacter baumannii* strain, AB5075-VUB. This compound, named HDC1, is a harmine-derivative previously designed to have anticancer properties. The anticancer screen showed that the compound acts as a protein synthesis inhibitor at a concentration of 0.7 µM. ^14^ In our study, HDC1 was shown to have bactericidal activity on AB5075-VUB with an IC_50_ of 48.23 µM. The cytotoxicity of the compound in human cells can thus be expected to be significant. This limits the potential of HDC1 as a new antimicrobial without further modification of the molecule. However, as multidrug-, extensively drug- and pandrug-resistant *A. baumannii* strains are emerging and spreading, every option deems to be explored.

Due to the high diversity between *A. baumannii* isolates, one of the main hurdles in the discovery of new compounds is to find compounds capable of targeting the majority of the isolates. Here, we report a compound to have inhibitory activity on all the tested and recent clinical *A. baumannii* isolates. The strains were chosen to be as diverse as possible: originating from different Belgian hospitals, different patients, varying anatomical infection sites and varying antibiograms. ^19^ Our test shows various degrees of growth inhibition: from complete to intermediate to only slight inhibition. In addition, the 4 reference strains used in our study all show a high degree of sensitivity to HDC1. Taken together, this raises the following questions (i) what contributes to this difference in growth inhibition levels, (ii) why do more recent clinical isolates show a higher resistance to HDC1 while the less recent reference strains are sensitive, (iii) what could be the target(s) of HDC1 and (iv) why is not the whole AB5075-VUB bacterial population killed by HDC1 in the tested conditions, since a significant bactericidal effect is observed? The phase variation observed in AB5075 might be the answer to the last question, for which bacteria in different states might show different levels of resistance against HDC1. ^22^ Interestingly, a recent study showed that HDC1 inhibits the phosphoserine phosphatase of *Mycobacterium tuberculosis*, MtSerB2. ^23^ MtSerB2 catalyzes the last step in the L-serine biosynthetic pathway and is involved in immune evasion mechanisms of *M. tuberculosis*. ^23^ In AB5075-VUB, a homolog of MtSerB2 is present: AbSerB. This indicates a putative target of HDC1 in *A. baumannii*. A possible explanation for the different growth inhibition profiles of the clinical isolates could be the presence of mutations in the *serb* gene. However, in our study, no correlation could be found between mutations in *serb* and the growth inhibition profiles of the *A. baumannii* isolates. Interestingly, it has been shown for *M. tuberculosis* that certain compounds are more efficient at inhibiting growth of the bacterium itself, than inhibiting the enzyme only, suggesting different mechanisms of action or intracellular accumulation of the compounds. ^24^ Additional resistance mechanisms, potentially countering such effects, could explain the higher resistance to HDC1 of some clinical isolates. Nevertheless, HDC1 has broad activity on all the tested recent clinical *A. baumannii* isolates of our study, which prompts the further exploration of this compound and/or cognate putative target in *A. baumannii*.

## Conclusion

In conclusion, HDC1 is a potent harmine-derived compound with antibacterial activity identified using a multidrug-resistant *A. baumannii* strain, that also significantly inhibits the growth of diverse, recent, and extensively drug-resistant clinical *A. baumannii* isolates.

## Author Contributions

Drafting of the manuscript: AB and CV. Corrections of the manuscript: AB, MVG, ME, CP, PB, TDH, OD, JW, SB and CV. HDC1 production: ME and SB. Preliminary drug screen: TQ, CP and CV. Experiments: AB, CW and CV. Data analyses: AB, MVG, JW and CV.

## Acknowledgements

We are grateful to Pr. Tom Coenye for providing us the strain AB5075 and Dr. Suzana Salcedo for the strains ATCC19606, ATCC17978 and DSM30011, as well as for fruitful discussions. We would like to thank Pr. Pierre Bogaerts, Pr. Te-din Huang and Pr. Olivier Denis for providing the recent clinical isolates of *A. baumannii*. We also thank the members of Namedic (NARILIS-UNamur), in particular Pr. B. Masereel and L. Pochet for their research that allowed setting up the chemical library used in the present screening.

## Transparency declarations

None to declare.

## Supplementary data

**Table S1.** Relationship between the absorbance (OD_600nm_) and the colony forming units (CFUs) per ml for the strain AB5075-VUB. The values are shown for overnight cultures that were diluted for accurate absorbance measurements. Rep = Biological replicate. At an OD_600nm_=1, the CFU/ml corresponds to 3.2±0.3 10^8^ CFU/ml.

|  | Rep 1 | Rep 2 | Rep 3 | Average | SD |
| --- | --- | --- | --- | --- | --- |
| **$OD_{600}$** | 6.4 | 6.5 | 6.3 | 6.4 | 0.1 |
| **CFU/ml** | $1.92 \cdot 10^9$ | $2.06 \cdot 10^9$ | $2.23 \cdot 10^9$ | $2.07 \cdot 10^9$ | $1.55 \cdot 10^8$ |

